# Treatment of murine autoimmune myocarditis with a novel monoclonal antibody that targets multiple inflammatory pathways

**DOI:** 10.64898/2026.03.27.714891

**Authors:** Stefano Toldo, Dror Luger, Aimee Vozenilek, Antonio Abbate, Jazmin Kelly, Eleonora Mezzaroma, Cyndya A. Shibao, Marwa A. Abd-ElDayem, Paul Klenerman, Ron Waksman, Renu Virmani, Jennifer A. Maynard, David G. Harrison, Moshe Y. Flugelman, Stephen E. Epstein

## Abstract

Severe forms of inflammation-induced acute and chronic myocarditis have a poor prognosis. Promising therapeutic efforts focused on monoclonal antibodies (mAbs) inhibiting inflammation-inducing molecules. However, most mAbs target only one or a limited number of such molecules. Since inflammation involves multiple redundant pathways, we postulated that an mAb inhibiting multiple inflammatory pathways would be a potent therapeutic agent. We initially tested the commercially available anti-natural killer (NK) cell mAb (anti-NK1.1), which binds a receptor expressed on NK cells and depletes them. Since NK cells are key cellular orchestrators of inflammation, by reducing their number, we aimed to inhibit multiple inflammatory pathways. Our initial studies demonstrated that administration of this antibody significantly improved myocardial outcomes in mouse models of acute myocardial infarction and of heart failure. Since NK1.1 is not expressed in human cells, we built on these promising preclinical results by developing a novel mAb targeting CD160 on human NK cells for evaluation as an immunosuppressive therapy. We found that the anti-CD160 mAb depletes *both* murine and human NK cells. We also found that, while CD160^+^ cells were largely present in the NK population, they also occurred among CD8^+^ and γ/δ T cell subsets in human cells. Anti-CD160 therapy entirely prevented the deterioration of the myocardial function of mice with autoimmune-induced acute myocarditis. This outcome suggests our novel approach for inhibiting multiple inflammatory pathways may provide a potent strategy for improving outcomes of inflammation-driven myocarditis, as well as of other inflammation-driven diseases.

**Key Points:** *Question:* Can the depletion of CD160^+^ cells prevent autoimmune-induced myocarditis?

*Findings:* In this study we found that CD160 is expressed by mouse and human natural killer cells and other subtypes of cytotoxic T cells, and that a monoclonal antibody targeting CD160 depletes NK cells. In a preclinical model of experimental autoimmune myocarditis, administration of the anti-CD160 monoclonal antibody prevented myocardial dysfunction and systemic inflammation.

*Meaning:* Our results are compatible with the hypothesis that early autoimmune-induced myocardial dysfunction is promoted by CD160^+^ cells, which elevate inflammation-induced *circulating* factors (or factors released by *tissue-resident* cytotoxic immune cells) that cause myocardial dysfunction in the absence of myocardial necrosis or fibrosis, and further, that targeting CD160^+^cells with a mAb that depletes NK cells (and probably CD160 expressing cytotoxic T cells) entirely prevents the deterioration of myocardial function in such mice. This outcome suggests our novel approach for inhibiting multiple inflammatory pathways may provide a potent strategy for improving outcomes of inflammation-driven myocarditis, as well as of other inflammation-driven diseases.

## Introduction

Infections pose an existential threat to species’ survival, which has led, over millions of years, to the evolution of robust inflammatory responses combating the lethal effects of infection.^1^ This includes the evolution of major redundancies in inflammatory pathways, a necessary adaptation to evolutionary pressures requiring a potent response to the multiple and changing pathogens threatening species’ survival.^1^ However, although inflammation is key to species survival, it can become excessive or inappropriately prolonged, leading to diseases initiated or exacerbated by inflammation^2^.

Globally, myocarditis occurs in about 4 to 14 of 100,000 people presenting as either chest pain, acute heart failure (HF), life-threatening arrhythmias, or cardiogenic shock, and is a prominent cause of sudden death.^3^ The most common cause of myocarditis is viral infection.^3–5^ Although viruses can directly injure the myocardium, viral-triggered immune-mediated reactions appear to be the principal cause of cardiomyocyte injury.^3,6^ Immune/inflammation-mediated myocarditis also derives from increased use of immune checkpoint inhibitors (ICI).^7^ Although transformational in the treatment of cancer, the activated T cells can cause off-target effects and serious autoimmune complications. Myocarditis occurs in 1-2% of ICI-treated patients.^7^

The development of monoclonal antibodies (mAbs) that inhibit inflammatory pathways is an effective therapeutic strategy that ameliorates inflammation-driven diseases.^8^ However, each of the currently available FDA-approved mAbs target only a single or a few inflammatory pathways. If the concept that redundancy of inflammatory responses is vital for species survival is correct, it would be reasonable to conclude that anti-inflammatory interventions blocking a limited number of inflammatory pathways would have important therapeutic limitations.

In a series of studies on the potential of mesenchymal stem cells (MSCs) to improve myocardial dysfunction occurring in murine models of acute myocardial infarction (AMI) and ischemia-induced chronic HF, we previously found that MSCs improved this dysfunction in both models.^9^ We further determined that a major responsible mechanism was MSC-induced depletion of natural killer (NK) cells—which are orchestrators of a diverse array of inflammatory pathways.^9^ We therefore hypothesized that the myocardial dysfunction occurring after acute MI and late post AMI was not only caused by direct ischemia-induced myocardial necrosis and fibrosis; in fact, *inflammation* also appeared to play an important mechanistic role. Most importantly, its inhibition could actually *reverse* a significant part of the dysfunction.^9,10^

In our efforts to translate these observations into a novel, effective therapeutic strategy, we demonstrated that a commercially available mAb (anti-NK1.1; BioLegend) that binds to the NK cell target NK1.1 depletes NK cells.^9^ Moreover, in murine models of AMI and ischemic-HF, we found that administration of anti-NK1.1 improved myocardial dysfunction.^9,11^

As NK1.1 is not expressed in human cells, we developed a novel anti-human NK cell monoclonal antibody that binds to CD160 (IFT-100), a glycoprotein present on subsets of natural killer (NK) cells in mice and humans.^12^ As we found, IFT-100 depletes both murine and human NK cells. We therefore decided, as our primary aim, to determine whether targeting CD160 would, like NK1.1, improve outcomes in a myocardial disease caused or exacerbated by inflammation. Importantly, we realized that moving forward with CD160 as the therapeutic target was a gamble, as we are unaware of any published studies demonstrating that targeting NK cells with an anti-CD160 mAb depletes the cells or has therapeutic potential. The gamble paid off.

We also decided to determine whether the myocardial benefits of an anti-NK cell strategy we found in models of myocardial dysfunction induced by ischemia were, in fact, limited to ischemia-induced dysfunction. Consequently, the current investigation was designed to determine whether a mAb-mediated anti-CD160 strategy is effective therapeutically in a murine model of *experimental autoimmune-induced myocarditis* (EAM), with the intention that if this proved to be the case, this study would provide the impetus for humanizing the mAb and proceeding with more detailed analyses of mechanisms necessary for advancing with a clinical trial.

## Methods

### Ethics Statement

Animal care and handling were in accordance with the National Institute of Health Guide for the Care and Use of Laboratory Animals (No. 85-23, revised 2011). The Animal Care and Use Committee approved all protocols at Washington Hospital Center, Washington, DC.

### Choice of a mAb

Since we aimed to develop a mAb for clinical use, we validated and patented IFT-100 (Application No. 62/955,569), a unique anti-NK cell mAb that binds to CD160, a 27 kDa glycoprotein present largely on subsets of natural killer (NK) cells, γ/δ T cells, CD8+ T cells and some CD4+ T cells. The efficacy of the chimeric antibody was tested in mice and human cells through flow cytometry (Figure 1).

**Figure 1.**
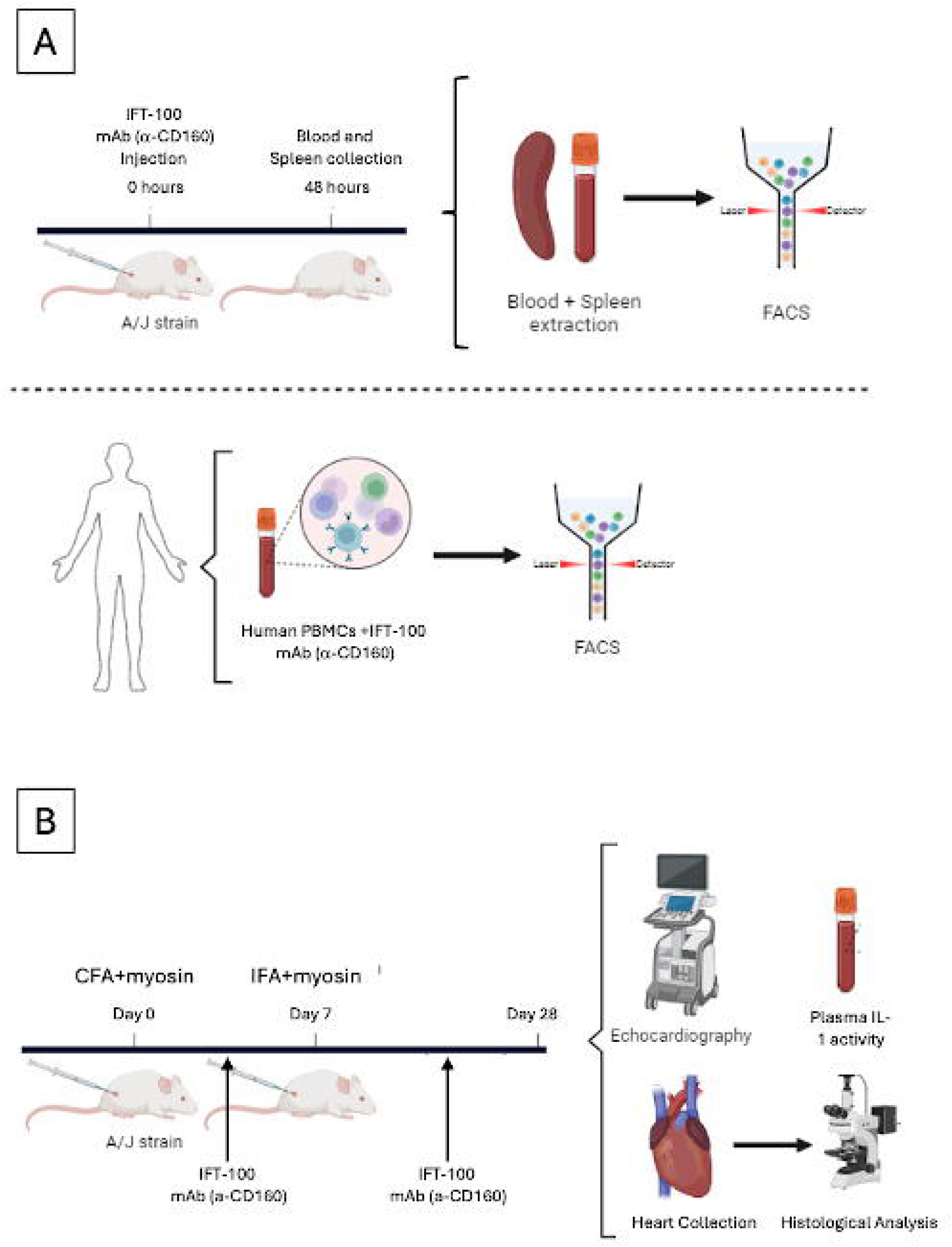
**Panel A:** Graphical representation depicting the experimental protocols and techniques utilized in the study. IFT-100, the chimeric anti-CD160 mAb, was injected into mice and, after 48 hours, the blood and the spleen were isolated to measure NK cell depletion. The effect of IFT-100 was also evaluated in human PBMCs. **Panel B:** Experimental protocol of murine autoimmune myocarditis. A/J mice were injected with Complete Freund Adjuvant and myosin on day 0, followed by a second injection with Incomplete Freund Adjuvant on day 7. IFT-100 was injected at day 2 and day 19 to deplete NK cells. Transthoracic echocardiography was performed at baseline and after 28 days. The hearts were collected to perform histological analysis. The plasma was used to measure Interleukin-1 activity. Abbreviations: IFT-100 (anti-CD160 monoclonal antibody); mAb (monoclonal antibody); PBMCs (peripheral blood mononuclear cells); FACS (fluorescence-activated cell sorting); CFA (Complete Freund Adjuvant); IFA (Incomplete Freund Adjuvant); IL-1 (Interleukin-1).

### Monoclonal Antibody Generation

Anti-human CD160 monoclonal antibodies were produced by GenScript (NJ, USA) via standard hybridoma technology. BALB/c mice were immunized with the target antigen, and splenocytes from high-titer animals were fused with SP2/0 myeloma cells. Hybridomas were screened by ELISA and flow cytometry for specificity to human CD160 and NK cells. Top-performing clones were subcloned, expanded, and purified using protein A/G chromatography. Final antibodies were validated for specificity and affinity, and cryopreserved for downstream applications.

### Chimeric Antibody Generation

A chimeric IgG1 antibody was developed by GenScript (NJ, USA) by fusing murine variable regions with human IgG1 constant domains. Variable heavy and variable light domain sequences from hybridoma cells were codon-optimized and cloned into cytomegalovirus-driven expression vectors. HEK293 cells were transiently transfected with plasmids encoding the heavy and light chains and cultured in serum-free medium. Antibody-containing supernatants were harvested and quantified by ELISA. Purification involved Protein A affinity chromatography followed by buffer exchange and size-exclusion chromatography to remove aggregates.

### In vitro depletion of CD160 cells from human peripheral mononuclear cells

Peripheral blood mononuclear cells (PBMCs) were isolated from healthy subjects, cultured in RPMI with or without IFT-100 (5 µg/ml) and human serum to induce the complement cascade. Cells were incubated overnight and underwent flow cytometry analysis to measure NK cell depletion from the PBMC pool.

### Flow cytometry

Mouse immune cells were isolated 48 hours after mice were treated with the IFT-100 mAb. Spleen cells and peripheral blood mononuclear cells were stained with antibodies against NKp46 and NK1.1 (Biolegend, San Diego, CA, USA) to define the specific NK cell population affected by the ITF-100 treatment. Human PBMCs were stained using antibodies against CD16 and CD56 (Biolegend). In all mouse and human experiments, cells were gated to exclude dead cells and to include singlets and CD45+ cells before NK cell measurements.

### Experimental autoimmune myocarditis (EAM) induction

EAM was induced in A/J mouse strain (Jackson, Bar Harbor, MA) using cardiac myosin (Genscript, NJ) as an autoimmune stimulator.^13^ Cardiac myosin was emulsified with Complete Freund’s Adjuvant (CFA) (at 1:1 ratio) and injected subcutaneously in two locations (100 μl/each) on the back of the mouse. Myocarditis was induced on day 0, and a boost of myosin emulsified with Incomplete Freund’s Adjuvant (IFA) was given on day 7 (Figure 1). Control mice were injected with CFA and IFA in absence of cardiac myosin. Mice receiving CFA+myosin were randomized to treatment with a control IgG (N=17, 50 mg/mouse) or IFT-100 (N=16, 50 mg/mouse). CFA-treated mice (controls, N=14) received an equal volume of vehicle (PBS). Treatments were administered on day 2 and 19 after myocarditis induction.

### Left Ventricular Function

Cardiac function was evaluated by transthoracic echocardiography using the Vevo 1100 (Visual Sonics, Ontario, Canada). Measurements were made a day before starting the experimental protocol, considered the baseline, and 28 days following the injection of myosin. Before each echocardiogram, mice were weighed and lightly anesthetized with sodium pentobarbital (30-50 mg/kg) to minimize distress. M-mode recordings of the midventricular section of the left ventricular (LV) were recorded in the parasternal short-axis view. Left Ventricle Ejection Fraction (EF) was calculated using the Teicholz formula. Cardiac output was assessed by multiplying stroke volume by heart rate. In addition, the LV long-axis fractional area change was calculated as the difference between the end-diastolic area and the end-systolic area measured through a longitudinal view of the LV. Echocardiography was performed and analyzed by an investigator who was unaware of the group allocation (study vs. control mice). At the end of the experiment (day 28), blood was drawn for cytokine analysis. Mice were euthanized, and hearts were harvested.

### Plasma Interleukin-1 activity

Interleukin-1 (IL-1) reporter cells (HEK-Blue™ IL-1 Receptor cells, Invivogen, San Diego, CA, USA) were used to measure the systemic activity associated with IL-1 in the plasma of mice.^14^ RPMI culture media (0.18 ml) was supplemented with 0.02 mL (10% final concentration) of EDTA plasma isolated from the mouse blood, with or without recombinant human IL-1Ra (1 µg), an antagonist of the IL-1Receptor, to measure and subtract the background signal. The cells were incubated for 24 hours before measuring reporter activity. The assay was repeated with three technical replicates for each sample and condition (+ or – IL-1Ra). IL-1 activity was defined as the optical density measured at 650 nm due to the SEAP Blue^TM^ (Invivogen) reagent conversion by reporter cells with the plasma, minus the activity of reporter cells with plasma and IL-1Ra, using a plate reader. In addition to the plasma of mice treated with CFA, plasma from control, naïve mice, was used to determine the effects of CFA treatment alone on IL-1 activity.

### Methods for histological examination of the mouse heart

Mouse whole hearts were fixed with 2% paraformaldehyde for 4-6 hours, dehydrated with a graded series of alcohol and xylene, and embedded with paraffin. The whole hearts were longitudinally cut into two parts to expose four chambers, placed in molds, and embedded in paraffin. Paraffin blocks were cut at 4 to 6 µm thickness and mounted in charged slides. The slides were stained with Hematoxylin and Eosin (H&E) and Masson’s Trichrome (MT). Pathological findings, including myocardial injury, inflammation, and fibrosis, were evaluated. Myocardial inflammation and fibrosis were scored using semi-quantitative scoring. Histology was performed and analyzed by an investigator unaware of group allocation (study vs. control mice). Inflammation and fibrosis were independently scored based on the extent and distribution of the specific staining. For inflammation, a score of 0 meant no inflammatory cell foci detected. A score of 1 was assigned to sections showing one small focal area of inflammation, a 2 to multiple small focal areas, and a 3 to diffuse areas of inflammation. For Fibrosis, a score of 0 was given to sections without fibrosis, a 1 to sections with one small focal area of collagen deposition, a two to multiple small focal areas, and a 3 to diffuse or multiple large areas of collagen deposition.

### Single-cell analysis of CD160^+^ cells

To examine the distribution of CD160 among immune cells in humans, we analyzed data from 4 healthy control subjects (mean age 29.5 ± 3.1 years, all female) who were included as controls in a larger study of Long COVID autonomic dysfunction. These control subjects were defined as individuals with or without confirmed SARS-CoV-2 infection who achieved complete recovery without any post-COVID sequelae, confirmed by clinical assessment for at least 3 months after infection or vaccination. Peripheral blood monocytes were barcoded for each subject and subjected to 10X sequencing using the Chromium platform. Data were analyzed in R using Seurat. After creating Seurat objects, low-quality cells and those with a mitochondrial content >10% were removed. Data were normalized and variable features identified. The threshold for clustering was identified using an elbow plot from the principal component analysis, and a Uniform Manifold Approximation and Projection (UMAP) was created. Cell clusters were identified using the canonical markers, CD3E (T cells), CD4 and CD8A (T cell subsets), NCAM1 and FCGR3A (NK cells), CD14 (monocytes), TRDC, TRGC1 and TRGC2 (γ/δ T cells), and CD160. CD160^+^ cells were first identified visually and then quantified as a percent of CD8^+^ T cells, NK and γ/δ T cells using Seurat. Comparison of genes expressed in CD160^+^ vs CD160^-^cells was performed in Seurat using the Wilcoxon Rank Sum test with the Benjamini-Hochberg correction for multiple comparisons, and the results were reported as the False Discovery Rate (FDR). Corrected FDRs < 0.05 were considered significant.

### Statistical analysis

Normal distribution of the values was assessed using the Kolmogorov-Smirnov test for normality. For non-normally distributed values, comparisons between multiple groups were performed using the Kruskal–Wallis test followed by Dunn’s test for multiple comparisons. Continuous variables are expressed as median and interquartile ranges. Data with normal distribution were analyzed using ANOVA and were expressed as the mean and standard deviation of the mean (SEM). Discrete variables are presented as percentages. Linear regression was used to model the relationship between variables. P values <0.05 were considered statistically significant.

## Results

### Anti-CD160 mAb recognizes and depletes *mouse* and human NK cells

After 48 h of IFT-100 IV injection in mice, splenocytes and PBMCs were analyzed through flow cytometry and showed a four-fold reduction of the NK cell population in the IFT-100 groups vs the control group in the spleen and a ten-fold reduction in the peripheral blood (Figure 2A). Following overnight incubation with IFT-100 and human serum, human peripheral blood mononuclear cells (PBMCs) were analyzed by flow cytometry and showed a 70% decrease in the NK cell population, as measured by CD16 and CD56, compared with the control group (Figure 2B).

**Figure 2.**
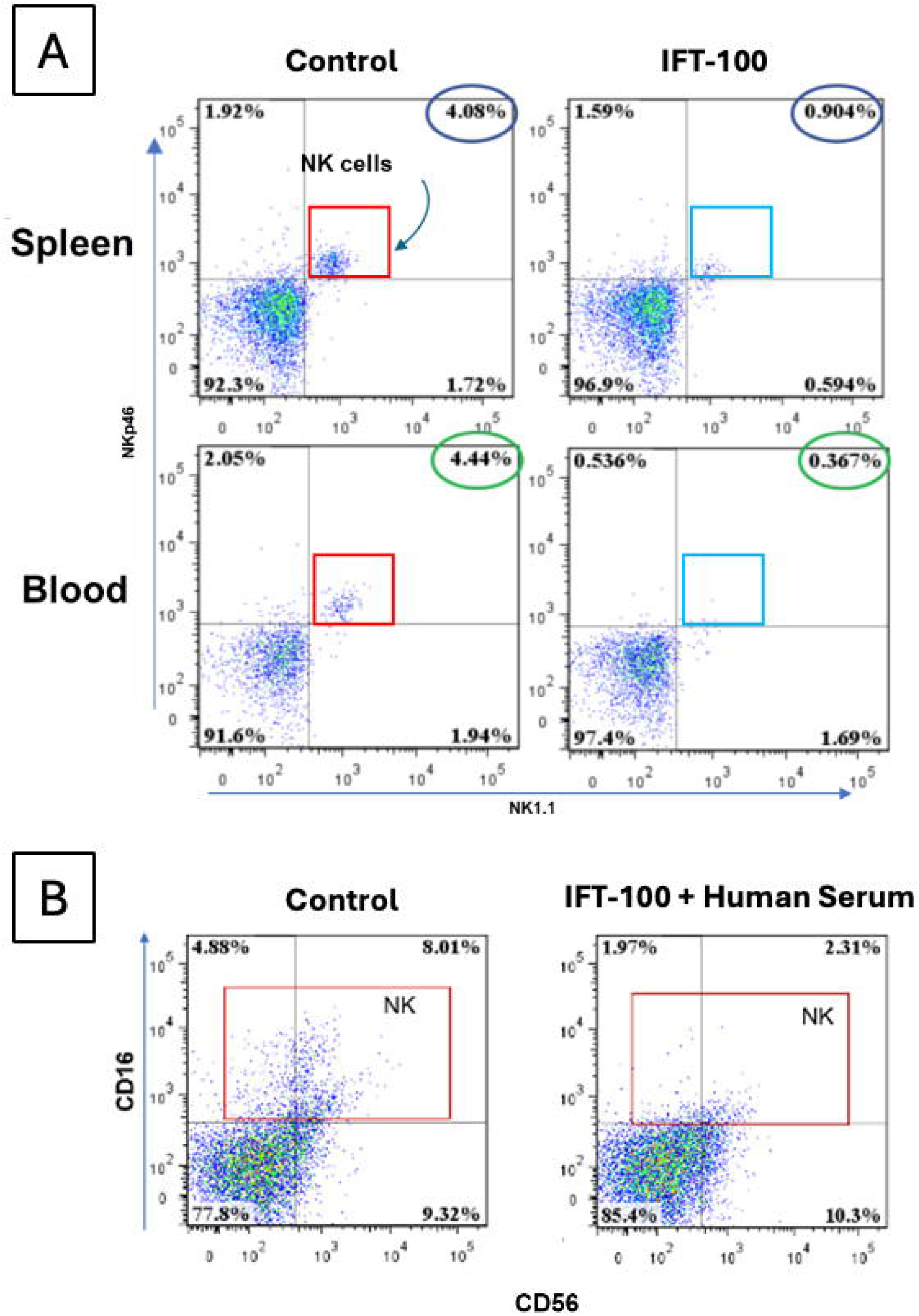
**Panel A:** Flow cytometry analysis showing that the anti-CD160 mAb (IFT-100) recognizes and depletes *mouse* NK cells in vivo in the spleen and in the blood. IFT100 was administered IV to mice. Splenocytes and blood PBMCs were analyzed 48h post-injection by flow cytometry. The boxes depict the NK cell population measured, the circles show NK cell percentages. **Panel B:** Flow cytometry analysis showing that the anti-CD160 mAb led to a 70% (8.01% to 2.31%) decrease in human NK cells. Human PBMCs were incubated with the mAb (plus human serum for complement activity). The boxes depict the NK cell population measured.

### Anti-CD160 mAb prevents the development of left ventricular dysfunction in mice with autoimmune-induced myocarditis, independent of inflammatory infiltrates and fibrosis

After 28 days, mice immunized against cardiac myosin but treated with the control IgG displayed significant reductions in LV ejection fraction (LVEF) when compared to the control mice treated with CFA (60.3±0.8% vs 53.7±1.7%; p<0.01, Figure 3A), cardiac output (10.3±0.7ml/min vs 7.7±0.4ml/min, p<0.01, Figure 3B), and area fractional shortening (37.0±2.5% vs of 28.7±1.7%, P<0.05, Figure 3C). Reductions in LVEF and cardiac output were not associated with a change in LV end-diastolic volume (Figure 3D). Treatment with anti-CD160 mAb in mice with induced myocarditis led to preservation of LVEF (61.9±1.3% vs 53.7±1.7%; p<0.01, Figure 3A), cardiac output (9.5±0.3ml/min vs 7.7±0.4ml/min, p<0.01, Figure 3B) and longitudinal area fractional shortening (37.3±1.5% vs 28.7±1.7%, P<0.01, Figure 3C). The hearts of mice were stained to assess fibrosis and inflammatory cell infiltration. The severity of the fibrotic and inflammatory responses during autoimmune induction varied considerably (Figure 3E, 3F, 3G). These anecdotal examples align with our group analyses. LVEF did not correlate with histological severity of either myocardial inflammation (Figure 3H), myocardial fibrosis (Figure 3I), or their combination (Figure 3J).

**Figure 3.**
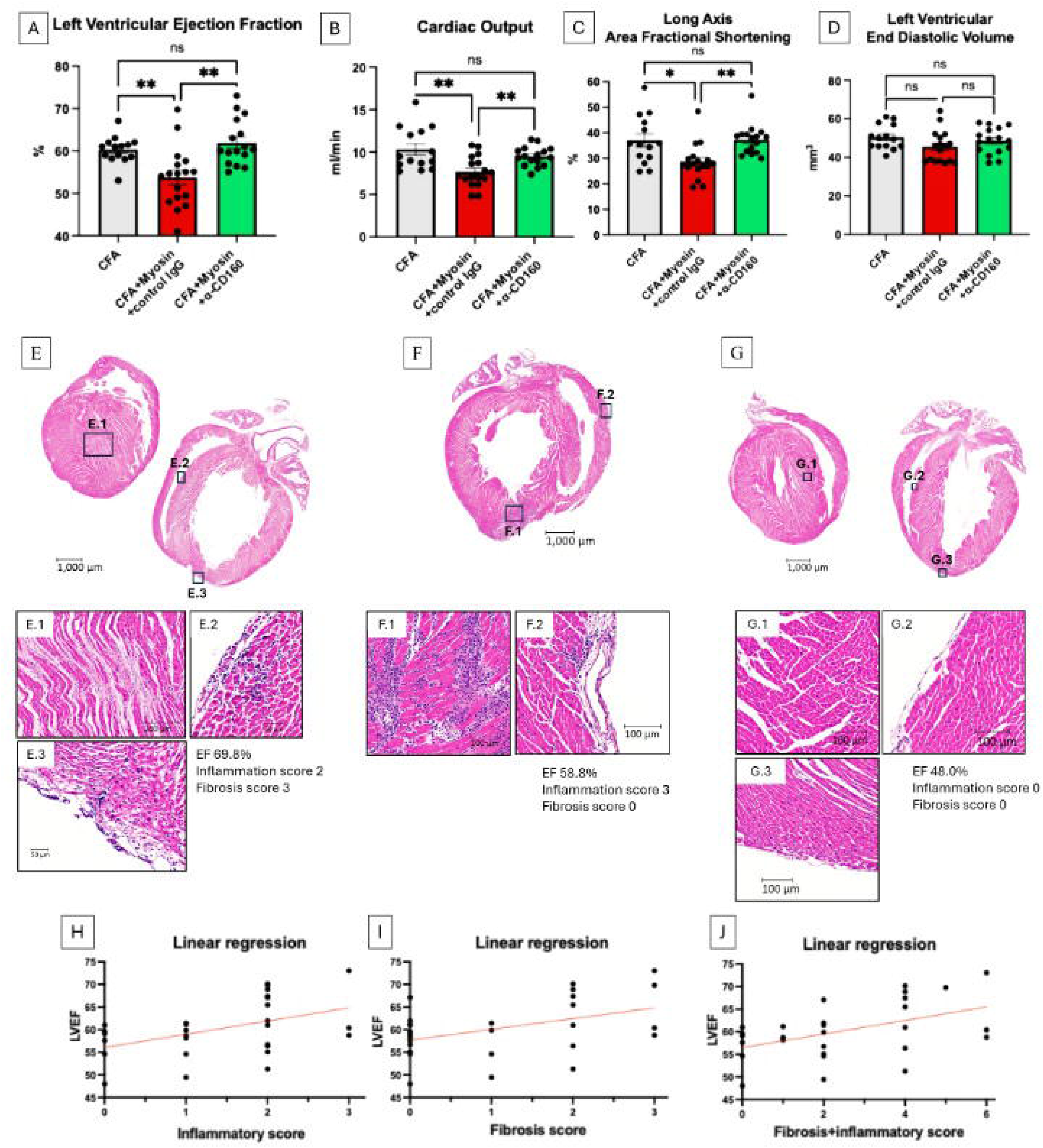
LVEF (**Panel A**), cardiac output (**Panel B**), longitudinal area fractional shortening (**Panel C**), and LV end-diastolic volume (**Panel D**) were measured 28 days after inducing autoimmune myocarditis, using transthoracic echocardiography. IFT100-induced depletion of NK cells prevented the development of LV dysfunction in mice with autoimmune myocarditis. *p<0.05; **p<0.01; ns = non-significant differences. Data represent mean ± SEM. **Panel E:** Representative histology images of the mouse heart with leukocyte infiltrates and diffuse fibrosis, and a high LVEF. E.1, E.2 and E.3 represent magnifications of areas of interest. **Panel F:** Representative histological image of a mouse heart with leukocyte infiltrates and diffuse fibrosis, and a normal LVEF. F.1. and F.2 represent magnifications of areas of interest. **Panel G:** Representative histological image of a mouse heart with no evidence of leukocyte infiltrates or fibrosis, and a lower LVEF. G.1, G.2 and G.3 represent magnifications of areas of interest. **Panels H, I, J:** Correlation analysis between LVEF, inflammatory infiltrates, and fibrosis. LVEF was correlated with the histological scores of myocardial inflammation (H), myocardial fibrosis (I), or their combination (J).

### Interleukin-1 plasma activity is decreased by CD160+ cell depletion

IL-1 activity results from the combined binding of the pro-inflammatory IL-1 isoforms, IL-1α and IL-1β, and the anti-inflammatory IL-1 receptor antagonist (IL-1Ra) on the IL-1Receptor (IL-1R)^15^. Elevated IL-1 activity is associated with LV dysfunction. Drugs that block IL-1, directly or indirectly, preserve LVEF^16^. In the present study, IL-1 activity was measured in plasma collected 28 days after the onset of the EAM. IL-1R reporter cells did not produce any signal when exposed to plasma from naïve untreated mice (Figure 4A). Plasma from mice treated with CFA alone elicited IL-1R activation, a not-unexpected result, as CFA is a pro-inflammatory primer. Plasma IL-1 activity was higher in mice with autoimmune myocarditis treated with control IgG. IFT-100 significantly reduced IL-1 activity in mice with autoimmune myocarditis (Figure 4A). Importantly, levels of IL-1 activity *inversely* correlated with LVEF (Figure 4B).

**Figure 4.**
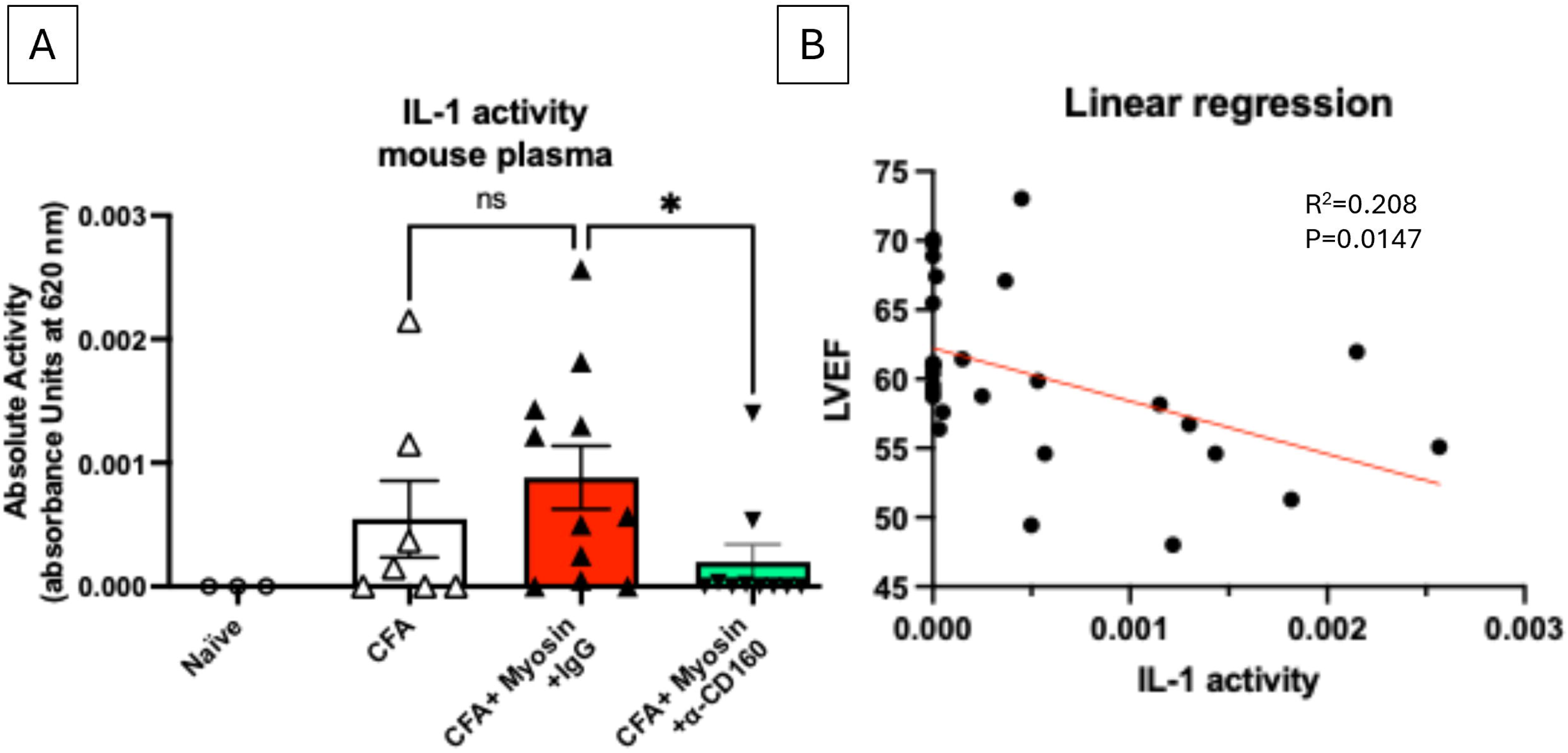
**Panel A:** IL-1 Receptor reporter activity measured in the plasma of naïve mice, control mice treated with CFA, and mice with autoimmune myocarditis treated with control IgG or IFT-100. IFT-100 reduced plasma IL-1 activity. *p<0.05. Data are presented as median and interquartile intervals. Circulating IL-1 activity tends to increase during autoimmune myocarditis, and administration of the anti-CD160 monoclonal antibody (mAb) significantly reduces IL-1 activity. **Panel B:** Group analysis correlation between the IL-1 activity measured in the plasma of mice (CFA, CFA+Myosin+IgG, CFA+Myosin+IFT-100) and their LVEF measured at 28 days after CFA treatment. IL-1 activity *inversely* correlated with the degree of impairment of LVEF.

### Single-cell sequencing to identify CD160^+^ cell phenotypes in human cells

To determine the distribution of CD160^+^ cells in humans, we performed single-cell sequencing on 4 human volunteers included in a larger study of autonomic dysfunction. CD160^+^ cells were present largely in the NK population (Figures 5A and 5B), but also among CD8^+^ T cells and γ/δ T cells (gamma delta T-cells). To gain insight into the functional consequences of CD160 expression in these cells, we performed an unbiased comparison of gene expression in CD160^+^ vs CD160^-^ cells. Figure 5C shows that after correction, there were 28 genes significantly higher in the CD160^+^ subgroup, and gene ontology analysis (panel 5D) showed these followed a pattern of promoting cell death and enhanced immune function vs. cells lacking CD160.

**Figure 5.**
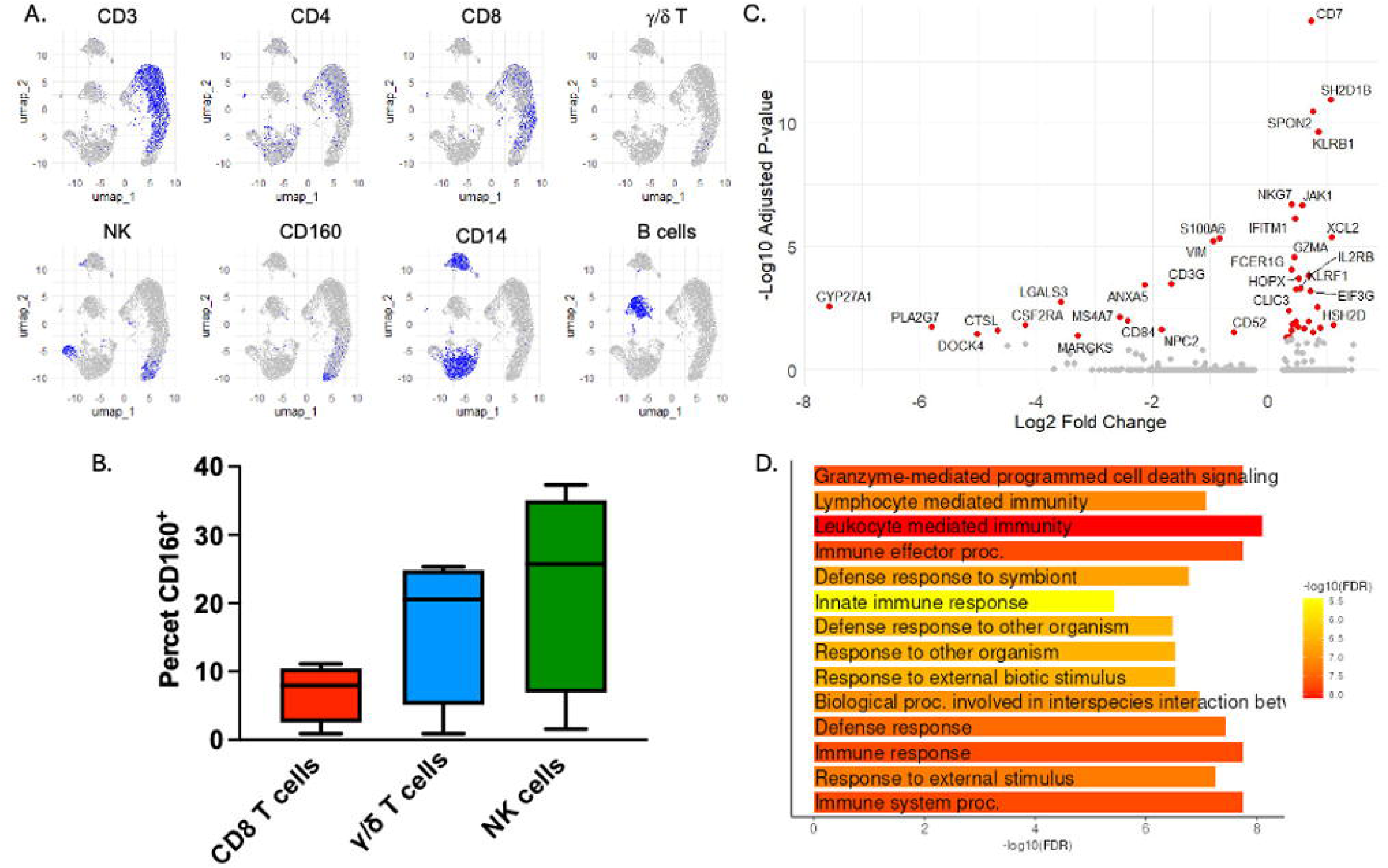
Single-cell sequencing of human peripheral blood mononuclear cells. **Panel A:** shows UMAPs of pooled cells from 4 normal volunteers, highlighting major leukocyte subsets, and including CD160^+^ cells. **Panel B:** shows the percentage of CD8^+^ T cells, g/d T cells and NK cells that express CD160^+^ cells. **Panel C**: volcano plot comparing genes expressed in CD160^+^ cells vs CD160^-^ cells. **Panel D:** gene ontology analysis of biological processes predicted to be affected by the overexpressed constellation of genes in CD160^+^ cells (from Shineygo 0.85 (https://bioinformatics.sdstate.edu/go/).

## Discussion

The reported results confirm the validity of our hypothesis: binding of CD160 by our novel mAb depletes NK cells and, most importantly, prevents myocardial dysfunction caused by autoimmune-induced myocarditis (Figure 3).

We have a long-standing interest in defining the role of inflammation as a cause of myocardial dysfunction in various clinical settings and in developing strategies that yield clinical improvement.^9,10,17–22^ Our current effort has been driven by the concept that there are major redundancies in inflammatory responses, accounting for the limited success of most current targeted therapeutic approaches, which mainly inhibit a limited number of such pathways.^23,24^ In initial studies, we depleted NK cells (which orchestrate numerous inflammatory pathways) using a commercially available mAb that binds to NK1.1 and depletes NK cells. Myocardial outcomes were significantly improved by mAb administration in mouse models of acute myocardial infarction and of chronic HF.

NK1.1 is not expressed in human cells. Since we are dedicated to developing a potent anti-inflammatory strategy as a *therapeutic* agent, we developed a novel anti-human NK cell monoclonal antibody that binds to CD160 (IFT-100), a 27 kDa glycoprotein present on subsets of NK cells and subtypes of cytotoxic immune cells in mice and humans.^12^

CD160 is one of the key receptors that modulate many of these cells’ activities.^12,25–27^ It has a predominant activating role in NK cells, enhancing cytotoxicity, cytokine production, proliferation, and migration; notably, inflammation also increases its expression on NK cells.^12,25–27^ A major role of CD160^+^ cells is to detect “non-self” peptides presented on class I major histocompatibility complexes.^28^ Upon such an encounter, CD160 recruits adapter proteins and signals cytotoxicity.^11^ Thus, CD160 has a major role in innate immune defences.^12,28,29^

To the best of our knowledge, there are no published studies demonstrating that targeting NK cells with an anti-CD160 monoclonal antibody leads to NK cell depletion in vivo. Nor have there been reports demonstrating that targeting CD160 has therapeutic potential. Existing work uses CD160 primarily as a marker and functional receptor, not as a validated NK-depleting target. We therefore conducted preliminary studies for the present investigation and found that IFT-100 depletes both murine and human NK cells. This presented a powerful opportunity for us to determine whether targeting CD160 with a mAb would, like NK1.1, improve outcomes of myocardial dysfunction caused or exacerbated by inflammation. To broaden the potential disease applicability beyond ischemia-induced dysfunction, we decided to determine whether the benefits of an anti-NK cell strategy are effective in a murine model of experimental autoimmune-induced myocarditis (EAM).

We performed single-cell sequencing of human PBMCs and confirmed that CD160 exists not only in NK cells, but also in CD8^+^ T cells, γ/δ T cells, and some CD4+ T cells. The gene expression profile in these CD160^+^ cells was consistent with CD160’s role in promoting cytotoxicity and pro-inflammatory responses. Thus, depletion of a broad array of CD160^+^ cells would likely reduce all these cell populations and thereby reduce their contribution to cytotoxicity and inflammation.

Recently, Tong et al. identified a striking expansion of CD8^+^ T cells in humans with acute myocarditis.^30^ These cells exhibited enhanced CD57 (B3GAT1) expression.^30^ Although NK cells were excluded from analysis, CD57 is also commonly expressed on subsets of NK cells.^31,32^ In CD8^+^ T cells, it is associated with T cell exhaustion and with increased cytotoxicity.^31,33^ Our dataset of normal humans did not reveal large numbers of B3GAT1^+^ cells in either the NK, CD8^+^ or CD160^+^ populations (data not shown). However, these individuals did not exhibit signs of acute disease. Nevertheless, these findings suggest that cytotoxicity plays a critical role in the pathogenesis of myocarditis and possibly other forms of HF.

Figure 6A depicts some of the multiple inflammatory pathways influenced by CD160-expressing cytotoxic immune cells, and shows how mAb-induced depletion of such cells can provide a potent therapeutic strategy for controlling inflammation-induced diseases.

**Figure 6.**
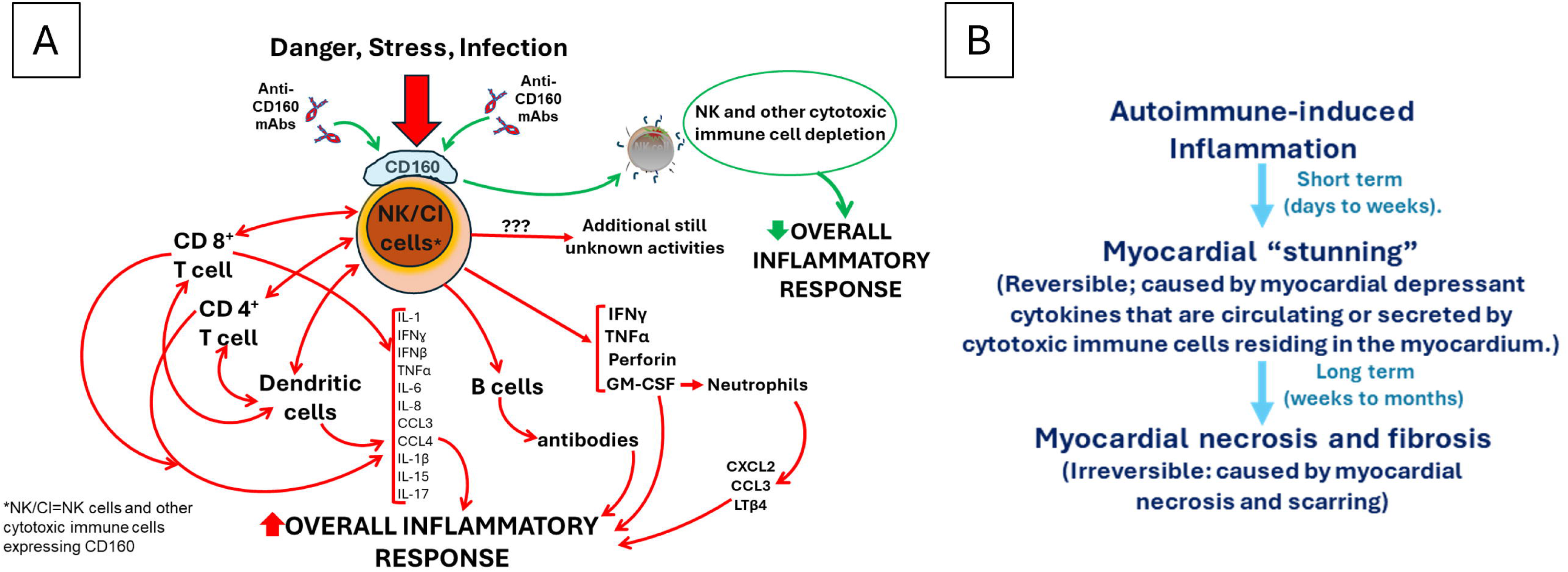
**Panel A:** Simplified schematic representation showing the possible direct and indirect effects NK cells could have on the inflammatory responses triggered by autoimmune-induced myocarditis—and thereby, the multiple pathways that could be influenced by an intervention depleting NK cells. This figure is hypothesis-generating rather than a demonstration of a proven pathway effect. Its major purpose is to demonstrate the redundancy of inflammatory responses, and the large number of inflammatory pathways NK cells influence by either direct or indirect effect. As noted, CD160 is also present on other cytotoxic immune cells. It is not clear at this point what the effects of an anti-CD160 mAb are on these cells and how important altering the activity of these cells is to the overall observed effects on inflammation. **Panel B:** Schematic illustration of the proposed mechanism leading to myocardial dysfunction in autoimmune-induced myocarditis. Various causal agents can lead to an autoimmune response that causes myocardial dysfunction. During the acute phase of autoimmune-induced myocarditis, myocardial dysfunction can be caused by the activities of inflammation-induced cytokines that are either circulating or derived from cytotoxic immune cells present in the myocardium. This type of dysfunction is likely reversible upon inhibition of the cytokines. The evolution from this reversible myocardial “stunning” to irreversible myocardial necrosis and fibrosis occurs over time.

### Mechanism responsible for impaired myocardial function

Our study also provides a novel hypothesis as to how myocardial dysfunction develops in autoimmune-induced myocarditis. We had expected to see extensive myocardial inflammation and fibrosis to explain the depressed myocardial function present in control mice.^13^ The lack of evidence of that occurring was unexpected. Semi-quantitative analysis revealed that the severity of impaired LVEF did not correlate with histological severity of either myocardial fibrosis, myocardial inflammation, or their combination, which were, in general, remarkably minimal (Figures 3H-J).

Therefore, it seems, at least in the model of autoimmune-induced myocardial dysfunction we used, and in its very early phase, myocardial dysfunction is not necessarily caused by inflammation-induced myocardial necrosis or extensive fibrosis formation. In fact, the early myocardial dysfunction seems to be caused in large part by inflammation-induced *circulating* factors (or factors released by *tissue-resident* cytotoxic immune cells) that can cause myocardial dysfunction *in the absence of myocardial necrosis or fibrosis —* something akin to myocardial stunning. Importantly, given the lack of myocardial necrosis and fibrosis, the mechanism leading to this myocardial dysfunction would, as discussed below, probably be reversible (Figure 6B).

Studies have shown that inflammatory cytokines can contribute to the development of myocardial dysfunction.^15,16,34^ In fact, IL-1β leads to a reversible type of myocardial dysfunction when administered to mice.^15,35^ Other studies further demonstrated that when IL-1 activity is blocked by a mAb in mice with HF, myocardial function improves.^36,37^ IL-1 blockade is also cardioprotective in mouse models of autoimmune or viral myocarditis^38,39^.

The present investigation strongly supports the concept that *circulating cytokines* contribute to the myocardial dysfunction seen early after the induction of the autoimmune-induced myocardial dysfunction. Thus, systemic IL-1 activity tends to increase during autoimmune myocarditis, and our mAb significantly reduces it (Figure 4). Importantly, the magnitude of decline in IL-1 activity correlates with the magnitude of improved myocardial function (Figure 4). We believe it is highly likely that the changes in IL-1 activity are also occurring with other cytokines, that the *combined* effects of multiple circulating cytokines contribute to myocardial dysfunction, and that our anti-CD160 mAb reduces not only IL-1 activity but also the detrimental myocardial effects of multiple circulating cytokines. Although we did not perform confirmatory studies on other cytokines, the findings presented here provide strong support for the validity of this hypothesis, as illustrated in Figure 6. Myocardial depressant cytokines also may derive from cytotoxic immune cells present in the myocardium.

*Our strategy to target cells rather than individual inflammatory cytokines* to control autoimmune-induced myocarditis is unique—but has important parallels in the evolving treatment of other autoimmune diseases. Thus, B cells play a role in the pathophysiology of autoimmunity, leading to the application of B-cell-depleting strategies to treat multiple diseases.^40^ Encouraging results have been reported in treating, for example, rheumatoid arthritis, systemic lupus erythematosus, and ulcerative colitis.^41,42^ We therefore believe that our targeting for depletion cytotoxic immune cells to treat autoimmune myocarditis—as well as other immune-driven diseases--is a reasonable strategy to continue developing.

Our study focused on the short-term outcome of autoimmune myocarditis and whether administering anti-CD160 mAb alters the outcome. It would not be too great a leap in hypothesis generation to postulate that the mAb-induced improved myocardial function evident early in the disease’s course may prevent the later development of irreversible components of myocardial damage—myocardial necrosis and fibrosis.

A *clinical* trial conducted by our group demonstrated the *clinical validity* of NK cell depletion for treating myocardial dysfunction. We examined the effects of MSCs on the myocardial dysfunction present in patients with chronic HF. A single infusion of MSCs decreased circulating levels of NK cells and significantly improved myocardial function. Importantly*, the magnitude of the MSC-induced decrease in NK cells significantly correlated with the magnitude of the increase in EF*.^9,10^ These data support the concept that inflammation importantly contributes to myocardial dysfunction in patients with chronic HF, and that this component of myocardial dysfunction *is reversible, meaning that when* this component is reduced by NK cell reduction, myocardial function improves.

A concern with any immune cell suppression therapy is the potential predisposition to infection.^43^ However, cytokines released by NK cells can suppress antiviral and antibacterial functions of CD8+ T cells.^44^ Interestingly, NK cell depletion in mice enhances clearance of persistent lymphocytic choriomeningitis virus and improves survival following intratracheal infection with Streptococcus Pneumoniae.^45,46^ Thus, NK cell depletion might actually prove useful in recovery from infection in some conditions.

While our study demonstrates that targeting and depleting CD160-expressing cells improves outcomes in autoimmune myocarditis, several limitations must be acknowledged. First, we focused on the effects of our mAb on NK cells, demonstrating it depletes these cells. However, we can only speculate on whether the mAb has functionally important effects on other cytotoxic immune cells that express CD160. Second, our mechanistic investigations relating to circulating cytokines were limited to changes in IL-1 activity; studies of additional cytokines and signaling pathways are needed to establish broader mechanistic depth. Third, the study lacks detailed immunohistologic analyses, which prevents us from establishing a potential role of cytotoxic immune cells residing in the myocardium. We also did not analyze CD160 Expression in NK and cytotoxic T cells in the context of disease.

Nevertheless, it is important to note that our research is the first to demonstrate the therapeutic potential of CD160 cell modulation in inflammation-driven myocardial injury; it introduces a novel immunomodulatory strategy capable of impacting multiple inflammatory pathways, not only by targeting NK cells but also other cytotoxic immune populations expressing CD160. The observed functional improvement in our murine myocarditis model, as well as the beneficial results we demonstrated in experimental AMI and chronic ischemia-induced HF, support the concept that CD160-directed immune intervention is a promising strategy for inflammation-driven myocardial disease. Finally, as inflammation leads to the initiation and/or progression of many other diseases, our results may provide a new and effective strategy for improving outcomes of the many other inflammation-driven diseases.

